# Single-Neuron Gene Expression Analysis Using the Maxwell® 16 LEV System in the Neural Systems and Behavior Course

**DOI:** 10.1101/107342

**Authors:** Rayna M. Harris, Adriane G. Otopalik, Colin J. Smith, Dirk Bucher, Jorge Golowasch, Hans A. Hofmann

## Abstract

Gene expression analysis from single cells has become increasingly prominent
across biological disciplines; thus, it is important to train students in these approaches. Here, we present an experimental and analysis pipeline that we developed for the Neural Systems & Behavior (NS&B) course at Marine Biological Laboratory. Our approach used the Maxwell^®^ 16 LEV simplyRNA Tissue Kit and GoTaq^®^ 2-Step RT-qPCR System for gene expression analysis from single neurons of the crustacean stomatogastric ganglion, a model system to study the generation of rhythmic motor patterns. We used double-stranded RNA to knockdown expression of a putative neuromodulator-activated sodium channel. We then examined the electrophysiological responses to known neuromodulators and confirmed that the response was reduced. Finally, we measured how mRNA levels of several ion channel genes changed in response. Our results provide new insights into the neural mechanisms underlying the generation and modulation of rhythmic motor patterns.

## INTRODUCTION

The Neural Systems & Behavior (NS&B) course at Marine Biological Laboratory (MBL) has provided intensive training in the concepts and methodology of behavioral neurobiology and systems neuroscience to doctoral and postdoctoral students since 1978. NS&B offers multiple training opportunities in modern approaches to the study of neural systems and behavior for the next generation of behavioral neuroscientists during early stages of their research careers. This approach includes intensive lectures and discussion, one-on-one interaction with scientists and extensive hands-on laboratory training with a variety of invertebrate and vertebrate preparations using state-of-the-art techniques and equipment. Building on this success, we aimed to enhance and expand the course by incorporating single-cell molecular and genomic techniques to complement and extend the electrophysiological characterization of specific neuronal subclasses, including identification of neurons in invertebrate preparations.

The stomatogastric nervous system of the Jonah crab, *Cancer borealis*, is a model system to study neural mechanisms underlying the generation of rhythmic motor patterns and their modulation (1). The crab stomatogastric ganglion (STG) contains just 26 neurons, and the electrical properties, functional connectivity and neuromodulatory regulation of this neural network have been well characterized. Thus, the STG is well suited for analysis of gene expression at the single-neuron level (2).

Of particular interest is the examination of genes encoding ion channels that regulate sodium and potassium conductances, especially channels regulated by neuromodulatory substances. We investigated whether a qPCR approach could be implemented in the NS&B course to quantify gene expression levels of ion channels that are important for regulating electrophysiology and behavior in this system

## METHODS

Adult Jonah crabs, *Cancer borealis*, were obtained from Yankee Lobster (Boston, MA) and maintained in tanks with running/ chilled seawater until used. Crabs were anesthetized, and the complete stomatogastric nervous system was dissected out of the animal and pinned out in a dish containing chilled (12–13°C) physiological saline solution. The ganglia were then desheathed, and the lateral pyloric (LP) and pyloric dilator (PD) neurons were identified based on morphology and electrical activity. One of the two PD and the LP neurons were injected with double-stranded RNA (dsRNA), targeting the sodium leak channel nonselective (*NaLCN*). After 24 or 48 hours, electrical activity of individual neurons and the entire network in response to bath-applied neuromodulators was examined before treating with collagenase and harvesting single PD and LP neurons using fine forceps under a dissecting scope. The PD and LP neurons were then individually placed in 200μl of homogenization buffer with 1-thioglycerol, frozen in liquid nitrogen and stored at –80°C until further processing (2–3 days later).

RNA was isolated using the Maxwell® 16 LEV simplyRNA Tissue Kit (Cat.# AS1280) according to the manufacturer's instructions and subsequently eluted with 30μl of nuclease-free water. A 9.5μl aliquot was reverse transcribed in a 20μl reaction using the GoTaq^®^ 2-Step RT-qPCR System (Cat.# A6010). Briefly, RNA, random hexamers and oligo(dT) were denatured at 70°C for 5 minutes, then reduced to 4°C. The tube was placed on ice, GoScript^™^ Reverse Transcriptase added and cDNA was synthesized on the Veriti^®^ Thermal Cycler (Applied Biosystems) with the following settings: 25°C, 5 minutes; 42°C, 60 minutes; 70°C, 15 minutes. The cDNA was diluted 1:5 in nuclease-free water, and 6μl was amplified in a 20μl real-time PCR using the GoTaq^®^ qPCR Master Mix and 0.25μM (final concentration) of each forward and reverse primer with the StepOnePlus^™^ Real-Time PCR System (Applied Biosystems). qPCR was performed using primers to amplify18S rRNA and three candidate ion channel genes: *shal*, *para* and *NaLCN*. The sequences of primers were as follows: 18S forward, AGGTTATGCGCCTACAATGG; 18S reverse, GCTGCCTTCCTTAGATGTGG; *shal* forward, CTACATCGGTCTTGGCATCA; *shal* reverse, AGATCCTGAACACGCGAAAC; *para* forward, TCGGTATGGTGCTGAAGGAT; *para* reverse, CAGTGTTCGGATACCCTTGG; *NaLCN* forward, ATGCTGACTGTGGGTGTGTC; *NaLCN* reverse, GACAGTGCCAAAGAGGATG.

Primer efficiencies were determined by amplifying a series of twofold dilutions of *C. borealis* whole ganglia cDNA covering four orders of magnitude of template (100–0.01ng of cDNA per reaction). The primer-specific amplification efficiency E (the amplification factor per PCR cycle) was derived from the slope of the regression using formula E=2–(1/slope). The results were plotted as the cycle of quantification (Cq) vs. log2[cDNA] (Figure 1). For all samples, the Cq values generated from the Applied BioSystems StepOnePlus^™^ software were converted to counts using count = E(Cq1–Cq), where Cq1 is the Cq of a single target molecule (set to 37 for all genes) using the R package MCMC.qpcr (3). Then a Markov Chain Monte Carlo (MCMC) algorithm was used to sample from the joint posterior distribution over all model parameters, thereby estimating the effects of all factors (cell type and dsRNA treatment) on the expression of every gene (3). The estimated stability of 18S rRNA was directly incorporated into the model as a control gene. To examine Pearson correlations in expression level between gene, counts were log transformed expression graphed using R (4).

**Figure 1.**
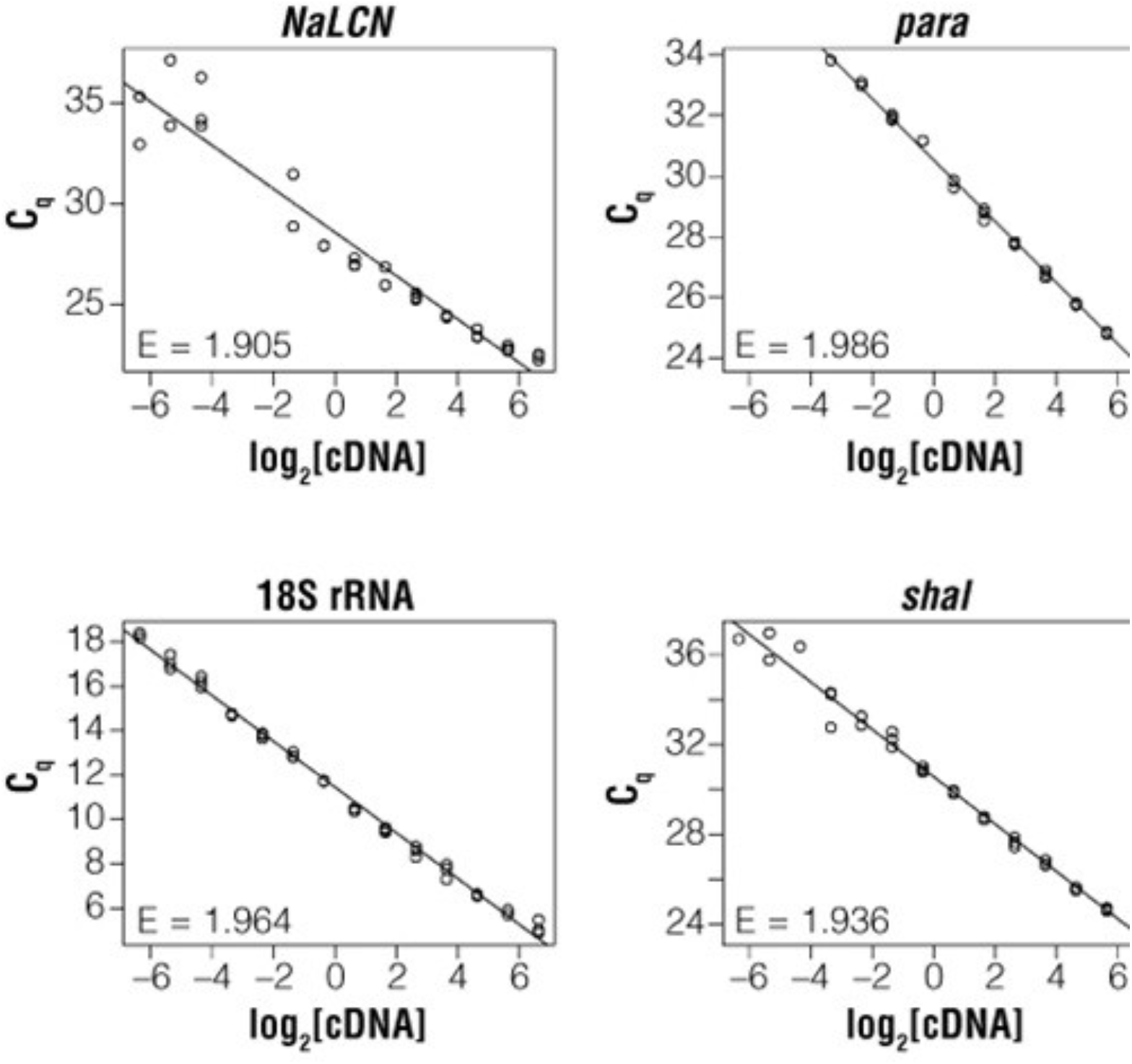
Primer efficiencies. Primer efficiencies (E, the amplification factor per PCR cycle) were determined by amplifying a series of twofold dilutions of *Cancer borealis* cDNA covering four orders of magnitude of template amount (100–0.01ng cDNA per reaction). The results were plotted as the cycle of quantification (Cq) vs. log2[cDNA]. E was derived from the slope of the regression using formula E=2–(1/slope). *NaLCN*: sodium leak channel nonselective; *para*: voltage-gated sodium channel; *shal*: potassium channel; 18S RNA: 18S ribosomal RNA subunit.

## RESULTS

Primer efficiencies (E) were all within the acceptable range (1.9–2.1; Figure 1). Sufficient RNA quantities were isolated from all samples for quantification of the four genes. A statistical analysis of changes in expression due to the factors of cell type (LP or PD) and treatment (dsRNA *NaLCN* treatment versus control) revealed that *NaLCN* expression is significantly decreased with dsRNA treatment independent of cell type (**p** = 0.048; Figure 2), but not for *para* (**p** = 0.992), *shal* (**p** = 0.784) or 18S RNA (**p** = 0.316). Across samples, *shal* and *para* expression levels were significantly correlated (r2 = 0.93, **p** = 1.39 × 10–5; Figure 3). Electrophysiological responses to two neuromodulators suspected of activating NaLCN channels (Crustacean cardioactive peptide [CCAP] and proctolin) were significantly reduced as expected if *NaLCN* expression was inhibited by dsRNA (data not shown).

## DISCUSSION AND CONCLUSIONS

Our results show that high-quality RNA can be obtained from single neurons in a course setting. Importantly, the amplification reactions produced valid and biologically meaningful data. *NaLCN* dsRNA treatment was sufficient at reducing levels of *NaLCN* RNA in the cells after 24 or 48 hours. The high degree of variance in expression between samples as seen in Figure 2 may be due to compensatory actions of the STG network through gap junctions and other molecular events. While there was no significant effect of dsRNA treatment on *para* or *shal* expression, we found that the expression of these two genes was highly correlated, indicating that the expression of these two ion channel-encoding genes may be tightly coregulated. We also showed that RNA interference in conjunction with electrophysiological recordings and manipulations provide an effective means for studying central pattern generation. Our results serve as a proof of concept that single-cell molecular approaches can be incorporated into the NS&B curriculum for generating insight into the neuromolecular mechanisms regulating rhythmic motor patterns in the crab STG.

**Figure 2.**
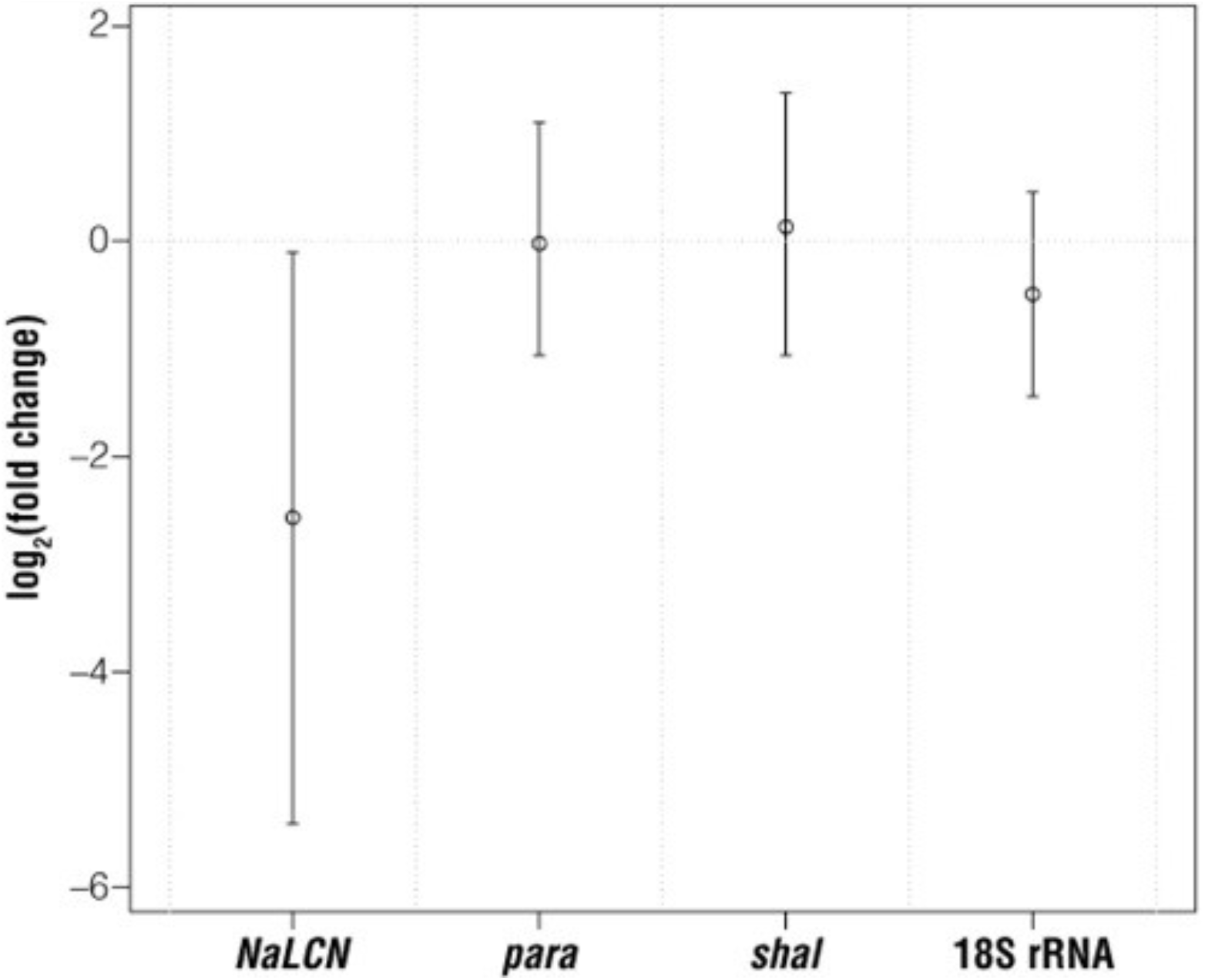
NaLCN dsRNA treatment reduces NaLCN expression. The effect of *NaLCN* dsRNA treatment on expression is plotted for each gene as log2(fold change), with positive and negative values indicating increased or decreased expression, respectively. The treatment occurred over 24 or 48 hours with both times included in the plotted data. The points are posterior means, the whiskers denote 95% confidence intervals.

**Figure 3.**
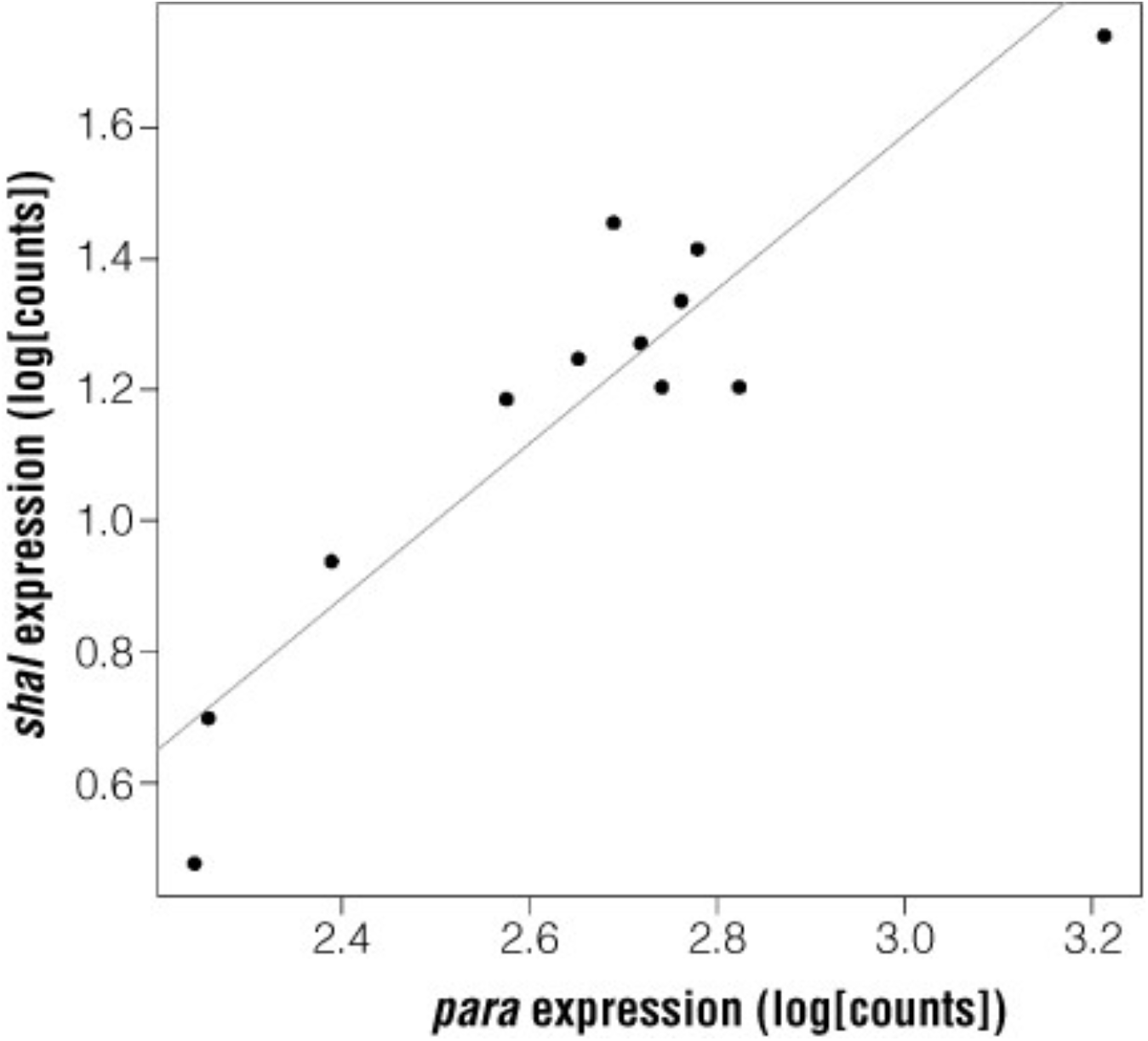
*para* and *shal* expression levels are correlated. A Pearson correlation of gene expression (log[counts]) for all samples analyzed shows that *para* and *shal* expression levels are significantly correlated (r2 = 0.93, p = 1.39 × 10–5). *para*: voltage-gated sodium channel; *shal*: potassium channel.

## ACKNOWLEDGEMENTS

We thank Nelly Daur and Dalia Salloum for dissecting the STGs used in this experiment, David Schulz for providing the *NaLCN* qPCR primer sequences and Deborah Baro for generously supplying the *NaLCN* dsRNA. We thank Dee Czarniecki and Doug Wieczorek from Promega for advice and for providing the Maxwell^®^ 16 LEV Instrument and molecular reagents. Finally, we thank the faculty and students of the 2013 NS&B course and attendees of the 2013 Gordon Research Conference in Neuroethology for fruitful discussions.

GoTaq and Maxwell are registered trademarks of Promega Corporation. GoScript is a trademark of Promega Corporation. StepOnePlus is a trademark of Applied Biosystems. Verti is a registered trademark of Life Technologies, Inc.

## AUTHOR CONTRIBUTIONS

All authors conceived and designed the experiment. RMH, AGO, and CS conducted the experiments, RMH and HAH analyzed the data and wrote the paper. All authors approved the final version of the manuscript.

